# AlignmentViewer: Sequence Analysis of Large Protein Families

**DOI:** 10.1101/269720

**Authors:** Roc Reguant, Yevgeniy Antipin, Rob Sheridan, Augustin Luna, Chris Sander

## Abstract

**Summary:** AlignmentViewer is multiple sequence alignment viewer for protein families with flexible visualization, analysis tools and links to protein family databases. It is directly accessible in web browsers without the need for software installation, as it is implemented in JavaScript, and does not require an internet connection to function. It can handle protein families with tens of thousands of sequences and is particularly suitable for evolutionary coupling analysis, facilitating the computation of protein 3D structures and the detection of functionally constrained interactions.

**Availability and Implementation:** AlignmentViewer is open source software under the MIT license. The viewer is at http://alignmentviewer.org and the source code, documentation and issue tracking, for co-development, are at https://github.com/dfci/alignmentviewer

**Contact:**, reaches all authors

## 1 Introduction

Multiple Sequence Alignment (MSA) analysis (e.g., analysis of sequence patterns, subfamilies, specificity residues, evolutionary couplings) and visualization allows researchers to extract information and gain a better understanding of protein families. MSA is a basic step in many analysis workflows, including: protein (i) structure prediction (Marks *et al.*, 2011), (ii) structure detection in flexible (‘disordered’) domains (Toth-Petroczy *et al.*, 2016), (iii) function prediction (Tamames *et al.*, 1998) and (iv) intracellular localization (Goldberg *et al.*, 2014).

A number of tools exist for the visualization of protein MSAs. MView (Brown *et al.*, 1997) was one of the first online browser-based MSA viewers, with alignments formatted as an HTML document and additional information for each sequence. Jalview (Waterhouse *et al.*, 2009) is a desktop-based tool developed in Java that is accessible through websites using an embeddable applet, but unfortunately the technology for these applets are no longer supported in popular browsers, such as Chrome. Another desktop application, AliView (Larsson, 2014), has features such as sorting, viewing, removing, editing and merging sequences from large nucleotide sequence datasets. MSAViewer (Yachdav *et al.*, 2016) is an interactive MSA visualizer in JavaScript that implements basic features of viewing, scrolling and motif selection. AlignmentViewer complements these other MSA tools and provides these features: (i) in-browser and serverless execution, (ii) visualization of very large MSAs, (iii) exploration of conservation patterns, (iv) sequence filtering, (v) logo display, (vi) pairwise sequence identity map (vii) spring-directed 2D-layout of protein distribution in sequence space (planned) and (viii) display of top-ranked evolutionary couplings (Marks *et al.*, 2011) (planned).

## 2 Implementation and functionality

AlignmentViewer is a platform agnostic tool built in JavaScript. It runs inside web browsers that are standards-compliant such as Chrome, Safari or Firefox. AlignmentViewer was developed with the D3 library (d3js.org) to produce dynamic and interactive data visualizations, and jQuery to simplify client-side scripting. Performance (speed) for large alignments was a major consideration in its development. The tool is entirely client-based running inside a web browser without communicating with the server, except for initiation and, as needed, downloading example alignment datasets. This enables the researchers to quickly start executing a local copy for online or offline use. Hyperlinks for lookup in background databases, such as Uniprot or PFAM, are made directly from the client. The tool has been thoroughly tested with many large alignments. An alignment with, e.g., 50,000 sequences (about 13MB of memory) loads in the Safari browser within one minute. Next, we describe the main features of AlignmentViewer.

### 2.1 Multiple Sequence Alignment (MSA) View

#### Alignment details

The MSA View tab shows summary information about the alignment: number of sequences, conservation and gap counts for each position, a sequence logo, and the alignment in one letter code. As a default, columns with gaps in the reference sequence (first row) are omitted in order to facilitate visual focus on sequence patterns relative to a protein of interest and to avoid extremely gappy alignment views typical of many MSA presentations. The amino acids are colored using a conventional coloring scheme, adopted from Mview, based on amino acid properties to facilitate recognition of patterns (alternative coloring schemes can easily be added).

#### Sequence attributes and sorting

In this view, sequences in the alignment can be sorted using one of four different methods: (i) the original order provided by the user, (ii-iii) % sequence identity (fraction of reference sequence identical to a second sequence, or vice versa, with gaps not counted), and (iv) sequence weights or other attributes, such as alignment profiles scores (e.g., HMM bit scores), provided by the user. The user can also filter proteins by sequence identity relative to a reference sequence or by percentage of gaps.

#### Hyperlinks to external databases and export

If the input MSA contains Uniprot IDs, the identifiers are linked to UniProt pages and species are indicated. After processing the alignment, the user can choose to export the sequences and save them locally.

**Fig. 1.**
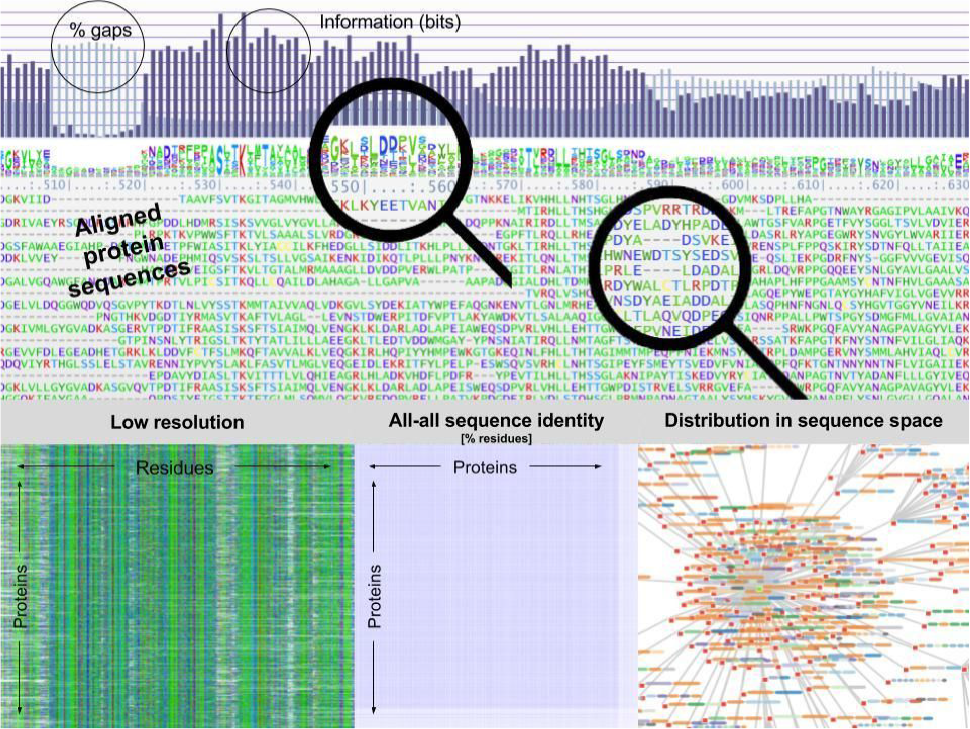
AlignmentViewer visualization of 14-3-3 protein domain family. Bar chart above the sequence alignment provides information on the sequence identity and percentage of gaps. The alignment consensus logo (just below the bar chart) summarizes the amino acid distribution at each sequence position. On the lower part of the figure, going from left to right we show the pixel view of the alignment, the sequence identity matrix, and a 2D distribution for the proteins (planned).

### 2.2 Pixel View Suitable for Large Families

To get on overview of the entire depth and breadth of an MSA we compress the amino acid letters into small rectangles of pixels, retaining the amino acid type coloring. The striking visual impression reveals patterns of conservation and variation. This is very useful to gain an intuitive view of sequence properties, noise at the uncertain edges of a protein family (‘twilight zone’), as well as subfamily distributions. The coloring scheme can be by (1) amino acid polarity, (2) by hydrophobicity (red to blue) or (3) by mutational difference (stronger color) in a sequence relative to the reference (first row) sequence. After any editing, one can save the bitmap representation of the alignment in an image file, e.g., for use in publications.

### 2.3 Distributions View

This tab shows statistical properties of the set of sequences in the alignment, including (i) sequence identity relative to the reference sequence, and (ii) min, max, and average sequence identity progression; (iii) a pairwise sequence identity matrix in which each pixel represents the degree of similarity between a protein on the x-axis against a protein on the y-axis, such that a block-diagonal structure of the matrix is indicative of distinct subfamilies; and (iv) a pairwise force-directed graph of the distribution in sequence space, which can be informative about phylogeny, uneven representation, as well as indicate suitability for evolutionary coupling analysis (planned).

### 2.4 Custom Data

Primary use cases for user-provided custom data (numerical attributes of a protein sequence) are analysis of sequence weights for evolutionary coupling analysis as well as comparison of different measures of sequence fit to the family, such as scores inferred from sequence profiles (e.g., PSSMs), Markov models or evolutionary Hamiltonian statistical energies. The sequences can be sorted by these scores.

### 3 Conclusion

AlignmentViewer is a lightweight online viewer for multiple sequence alignments in a wide size range with focus on usability and performance. The JavaScript frameworks chosen enables the use of the tool in many standards-compliant browsers. The architecture of AlignmentViewer allows its use without software installation and without an internet connection after loading. The visualization capabilities, analysis features and metrics in AlignmentViewer are useful in many areas of biology, especially evolutionary, structural, synthetic and chemical biology. Further enhancements are planned: visualization of species diversity, predicted contact maps, and organization by sequences subfamilies with specificity residues. Development of AlignmentViewer would benefit enormously from active engagement of interested members of the open source community

## 4 Acknowledgements

We thank Boris Reva and Debora Marks for constructive discussions. The project was supported by the Human Frontier Science Program (HFSP), the National Resource for Network Biology (NRNB), the Department of Cell Biology at HMS and the NIGMS.

